# The spliceosome inhibitors isoginkgetin and pladienolide B induce ATF3-dependent cell death

**DOI:** 10.1101/821363

**Authors:** Erin J. Vanzyl, Hadil Sayed, Alex B. Blackmore, Kayleigh R.C. Rick, Pasan Fernando, Bruce C. McKay

## Abstract

The spliceosome assembles on pre-mRNA in a stepwise manner through five successive pre-spliceosome complexes. The spliceosome functions to remove introns from pre-mRNAs to generate mature mRNAs that encode functional proteins. Many small molecule inhibitors of the spliceosome have been identified and they are cytotoxic. However, little is known about genetic determinants of cell sensitivity. Activating transcription factor 3 (ATF3) is a transcription factor that can stimulate apoptotic cell death in response to a variety of cellular stresses. Here, we used a genetic approach to determine if ATF3 was important in determining the sensitivity of mouse embryonic fibroblasts (MEFs) to two splicing inhibitors: pladienolide B (PB) and isoginkgetin (IGG), that target different pre-spliceosome complexes. Both compounds led to increased ATF3 expression and apoptosis in control MEFs while ATF3 null cells were significantly protected from the cytotoxic effects of these drugs. Similarly, ATF3 was induced in response to IGG and PB in the two human tumour cell lines tested while knockdown of ATF3 protected cells from both drugs. Taken together, ATF3 appears to contribute to the cytotoxicity elicited by these spliceosome inhibitors in both murine and human cells.

## Introduction

Most protein-coding genes are transcribed into a precursor RNA (pre-mRNA) that requires significant processing to produce mature mRNA. One of the more complex modifications is the removal of introns and ligation of exons through the process of pre-mRNA splicing [1]. The major spliceosome is responsible for the removal of the vast majority of introns, so it is usually referred to as the spliceosome[1]. The spliceosome is composed of five small nuclear RNAs (snRNAs), U1, U2, U4, U5 and U6, which are each associated with specific proteins yielding their corresponding small nuclear ribonucleoprotein complexes (snRNP). These snRNPs bind sequentially to *cis-*acting elements in pre-mRNA including the 5’ and 3’ splice sites, the branch sequence, and polypyrimidine tract, leading to the assembly of a functional spliceosome through the E, A, B, B* and C pre-spliceosome complexes [2].

A variety of small molecules interfere with the spliceosome at distinct points in assembly. PB is a spliceosome inhibitor derived from *Streptomyces platensis* that targets the SF3B1 protein [3]. This protein is a component of the U2 snRNP complex so PB prevents the formation of the A complex [3]. IGG is a natural compound derived from *Ginkgo biloba*, that affects the spliceosome at a subsequent step, preventing the transition from the A to the B complex by inhibiting the stable binding of the U4/U5/U6 tri-snRNP complex [[4]. These two compounds affect different stages of spliceosome formation and it is unclear if they induce cytotoxicity through one or more common pathways [5, 6].

Activating transcription factor 3 (ATF3) is a member of the ATF/CREB transcription factor family and is activated in response to a variety of stresses including DNA damage, inflammation, endoplasmic reticulum (ER) stress and oxidative stress [7-10]. ATF3 has been shown to have a variety of functions, one of which is regulating the expression of proapoptotic genes, such as GADD153/CHOP [11]. We sought to determine if ATF3 plays a role in determining the sensitivity of mammalian cells to the inhibition of pre-mRNA splicing. Here we report that ATF3 is induced in murine and human cells and that deletion of ATF3 in mouse embryonic fibroblasts (MEFs) and knockdown of ATF3 in HeLa cells protects them from the lethal effects of spliceosome inhibitors. This work provides important insight into the genetic basis of cell death induced by spliceosome dysfunction.

## Materials and Methods

### Cell culture and drug treatment

ATF3^+/+^ and ATF3^−/−^ MEFs were obtained from Dr. Jim Dimitroulakos (Ottawa Hospital Research Institute)[12] and grown in DMEM High Glucose media (Hyclone, San Angelo) supplemented with 10% fetal calf serum (FBS) (Gibco, ThermoFisher Scientific, Ottawa) and 90 units/ml penicillin, 90 ug/ml streptomycin (Hyclone). HCT116 and HeLa cells were obtained from the American Tissue Type Collection (Cat #: CCL-247 and CCL-2,, Manassas, VA) grown in McCoys and DMEM media (Hyclone), respectively, supplemented with a 3:1 mixture of Newborn Calf Serum (NBCS) (Gibco, Thermo Fisher Scientific) and Fetal Calf Serum (FBS) (Gibco, ThermoFisher Scientific), and 90 units/ml penicillin, 90 ug/ml streptomycin (Hyclone). Cell were seeded at a density of 500,000 cells/6 cm or 25,000 cells per well of a 96-well plate, 24 hours prior to treatment, depending on the assay. Cells were treated with up to 30 μM IGG (Calbiochem), up to 25nM PB (ChemCruz), 200nM actinomycin D (act D) (Sigma-Aldrich, St. Louis, MO), an equivalent volume of dimethyl sulfoxide (DMSO) as the vehicle control for IGG or left untreated.

### Reverse transcription (RT) and real time quantitative polymerase chain reaction (RT-qPCR)

RNA was isolated using the EZ-10 DNAaway RNA Miniprep Kit (Biobasics, Markham ON) and quantified using a DeNovix DS-11 Spectrophotometer (Wilmington, DE). Equal amounts of RNA were converted to cDNA using the High Capacity cDNA Reverse Transcription kit (ThermoFisher Scientific). For MEF experiments, ATF3 mRNA expression was determined using Bioline SensiFAST™ Probe HI-ROX Master Mix (FroggaBio Inc, Toronto, ON) with SYBR green. ATF3 expression was normalized to the average of two housekeeping genes; β-actin (ACTB) and Peptidylprolyl isomerase A (PPIA) using the following primers: ATF3 (F-AAGACAGAGTGCCTGCAGAA, R-GTGCCACCTCTGCTTAGCTC), ACTB (F-GATCTGGCACCACACCTTCT, R-GGGGTGTTGAAGGTCTCAAA) and PPIA (F-AGCTCTGAGCACTGGAGAGA, R-GCCAGGACCTGTATGCTTTA). Primers were synthesized by Integrated DNA Technologies (Coralville, Ia). For human samples, ATF3 mRNA expression was measured using the Hs00231069_m1 TaqMan^®^ Gene Expression Assays (ThermoFisher Scientific) with GAPDH serving as a sample control (Hs02758991_g1).

### Immunoblotting

Cell monolayers were rinsed with phosphate buffered saline (PBS) pH 7.6 and collected in 300μl of 1% SDS. Samples were sonicated (QSonic, Newton), and protein was quantified using the Bio-Rad Protein Assay (Bio-Rad, Philadelphia). Equal amounts of protein were denatured in NuPAGE™ LDS Sample Buffer and NuPAGE™ Sample Reducing Agent (10X) (ThermoFisher Scientific, Ottawa), heated for 10 minutes at 70°C and loaded onto NuPAGE 4-12% Bis-Tris gels (Fisher Scientific). Protein was then transferred to a nitrocellulose membrane (Bio-Rad), and transferred proteins were stained using 1mg/ml Ponceau S Red in 1% glacial acetic acid.

Membranes were blocked for 30-60 minutes in 5% Milk in Tris Buffered Saline Tween (TBST) pH 7.6 (Tris Buffered Saline, 0.1% Tween 20). Membranes were incubated overnight at 4°C in primary antibody, anti-ATF-3 (sc-188, Santa Cruz, CA) or anti-beta-actin (A2228, Sigma) diluted 1:200 for ATF3 or 1:1000 for Actin in 0.5% Milk in TBST pH 7.6. Membranes were washed 4×5 min in TBST, then incubated in secondary antibody (HRP-conjugated goat anti-mouse or goat anti-rabbit from Abcam) for 1-2 hours. Membranes were washed 4×5 min in TBST, then incubated for 5 minutes in 600μl of Clarity™ Western ECL substrate (Bio-Rad), and imaged using the Fusion FX5 gel documentation system (Vilber Lourmat, France).

### Flow cytometry

Floating and adherent cells were collected and combined 48 hours following treatment by centrifugation following trypsin treatment. The supernatant was removed, and the cell pellet was washed twice in PBS, fixed in 1ml ice cold 70% ethanol, and stored at −20°C for a minimum of 30 minutes. Cells were centrifuged, ethanol was then removed, and the pellet was washed twice in PBS and resuspended in 200μl of 20μM Propidium Iodide (PI) stain (Sigma-Aldrich) with 73μM RNaseA (Fisher Scientific). Red fluorescence was measured with a BD Accuri C6 flow cytometer and analyzed using the BD Accuri C6 software.

### Cell viability assays

ATF3^+/+^ and ATF3^−/−^ MEFs were seeded in a 96-well plate 24h hours prior to treatment for both cell viability assays. The trypan blue dye exclusion assay was used as an early indicator of cell death detected at 24 hours, prior to cell detachment. For trypan blue dye exclusion assays, cells were detached using trypsin 24 hours after treatment. Ten ul of trypan blue (0.4% in PBS) was added to each well of the 96-well plate. After two minutes, an aliquot of cells was transferred to Dual-Chamber cell counting slides (Bio-Rad, Hercules) and read using the Biorad TC20 Automated Cell Counter.

A luminescent ATP based cell viability assay was used as an indicator of a later stage of cell death after significant cell detachment was detected (48 hours). The CellTiter-Glo^®^ Luminescent Cell Viability Assay (Promega, Madison) was used according to manufacturer’s instructions to determine ATP content, to estimate cell viability. Briefly, 100 μl of media was removed from each well and mixed with an equal volume of CellTiter-Glo Reagent. After 2 minutes of shaking, the samples were incubated at room temperature for 10 minutes and read using the Cytation 5 96 well plate reader (BioTek, Vermont) and Gen5 software (version 2.07.17) at an integration time of 1.0 s per well.

### siRNA-mediated knockdown of ATF3

SMARTpool ON-TARGETplus siRNA (cat#: L-008663-00-0005 and D-001810-10-05, Dharmacon Inc, Chicago) were prepared in siRNA buffer as recommended by the manufacturer. HeLa cells were seeded at 2×10^6^ cells per 10 cm dish, in antibiotic free growth media 24 hours prior to transfection. The siRNAs were combined with lipofectamine 3000 as recommended by the manufacturer (ThermoFisher Scientific) to a final concentration of 50 nM in antibiotic and serum free media. The siRNAs were added to cells for 48 hours, after which transfected cells were seeded at 5×10^5^ cells per 6 cm dish for mRNA and protein collection and 1.25×10^5^ in 6 well dishes for flow cytometric analysis. Cell were treated 24 hours later and samples were collected at the indicated times for RNA, protein, and apoptosis assays from the same siRNA transfection.

## Results

### ATF3 is induced and contributes to cell sensitivity in mouse embryonic fibroblasts

Pre-mRNA splicing is essential for viability and pre-mRNA splicing inhibitors are cytotoxic through an unknown mechanism. It was recently reported that IGG-induces ATF3 indirectly through ATF4, however it was unclear if ATF3 contributed to cell death [13]. Therefore, we used a simple genetic approach to determine if ATF3 contributes to cell death following spliceosome inhibition using isogenic MEF strains that either expressed or did not express ATF3 (ATF3^+/+^ and ATF3^−/−^ MEFs, respectively). ATF3 mRNA levels increased significantly in the ATF3^+/+^ MEFs in response to PB but not IGG exposure (Fig 1A). ATF3 protein increased in response to both drugs, however we detected a greater increase in response to PB (Fig 1B). As expected, ATF3 protein was not detected in the ATF3^−/−^ MEFs (Figure 1B). We then assessed sensitivity to apoptosis by one parameter flow cytometric analysis of sub G_1_ DNA content in both cell lines following exposure to IGG and PB (Fig. 1B). Both drugs led to an increase in the proportion of cells with less than diploid DNA content, however PB led to a greater increase than IGG (Fig 1C). Importantly, cells deleted of ATF3 were significantly protected from drug-induced apoptosis compared to isogenic controls following exposure to both drugs. Therefore, ATF3 appears to contribute to PB-and IGG-induced apoptosis in MEFs.

**Fig 1.**
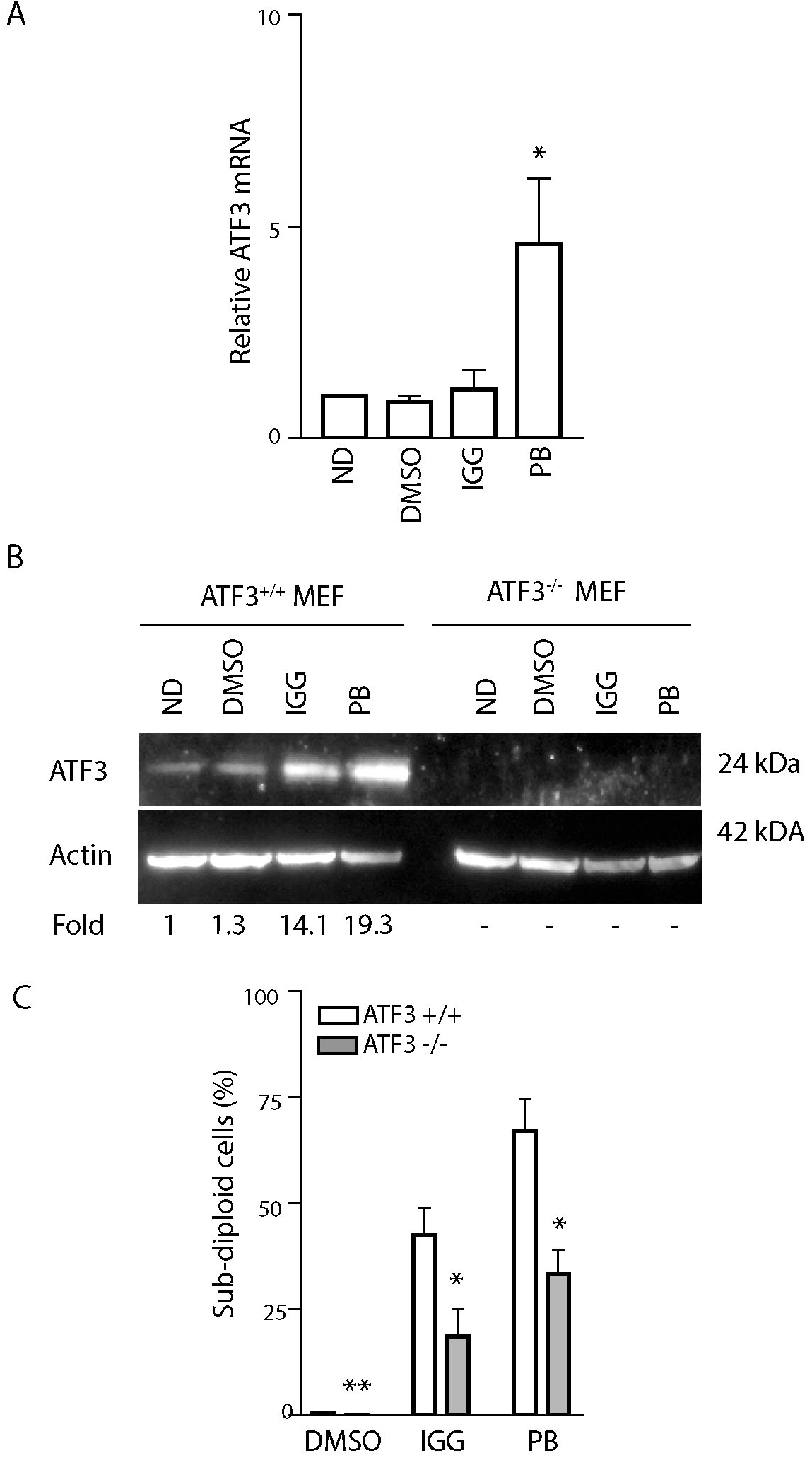
ATF3 is activated in response to IGG and PB treatment in MEFs. ATF3^+/+^ and ATF3^−/−^ MEFs were left untreated, or exposed to DMSO, 30μM IGG or 25nM PB for 24 hours and RNA and protein were collected (A and B). (A) ATF3 mRNA levels were quantified in 3 independent experiments in control MEFs using qRT-PCR, and statistical analysis was performed using a one-way ANOVA followed by Tukey’s post test. (B) Immunoblot analysis was performed with the indicated antibodies. Similar results were obtained in 4 independent experiments and the mean fold change is indicated below the blot. (C) Flow cytometric analysis of sub G1 content was performed 48 hours following treatment of ATF3^+/+^ and ATF3^−/−^ MEFs. Each value represents the mean, +/− SEM, of at least 3 independent experiments. The ‘*’ and ‘**’ indicate that the value is significantly different from that of its ATF3 expressing control (p<0.05 and p<0.01, respectively) by unpaired t-test.

The sensitivity of MEFs to spliceosome inhibition was further evaluated with two additional independent assays. First, ATF3^+/+^ and ATF3^−/−^ MEFs were exposed to either IGG or PB for 24h and cell viability was assessed by trypan blue dye exclusion (Fig 2A). Trypan blue is unable to penetrate viable cells with intact cell membranes, whereas non-viable cells that have lost membrane integrity take up the dye. Both splicing inhibitors (PB and IGG) decreased cell viability in ATF3^+/+^ MEFs. The surviving fractions of ATF3^−/−^ MEFs were significantly higher than parental controls following exposure to both drugs. In fact, there was no detectable decrease in viability in the ATF3^−/−^ MEFs with this assay.

**Fig 2.**
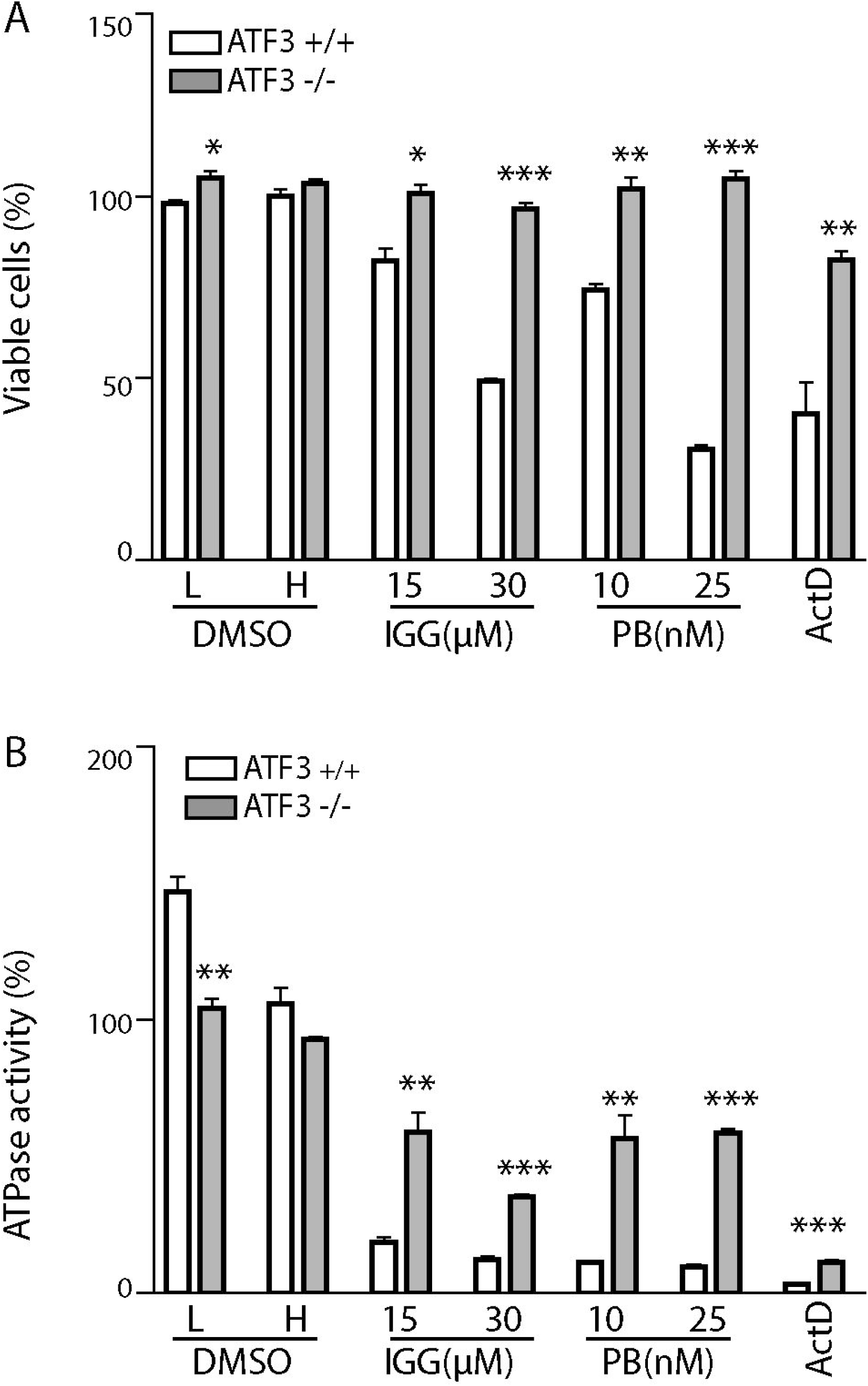
The absence of ATF3 in MEFs increases cell viability following PB and IGG treatment. ATF3^+/+^ MEFs (white) and ATF3^−/−^ MEFs (grey) were incubated with 30μM IGG, 25nM PB and 200nM act D for 24h (A) or 48h (B). L and H refer different volumes of DMSO required to serve as vehicle controls for the 2 concentrations of IGG. (A) A trypan blue exclusion assay was used to assess membrane integrity, as an indicator of cell viability. (B) The CellTiter-Glo^®^ Luminescent Cell Viability Assay was used to estimate metabolic activity as an indicator of cell viability. Each bar represents the mean of 3 independent experiments, +/− SEM. Statistical significance between cell lines for each treatment was completed using a t-test (*p<0.05, **p<0.01, ***p<0.001).

Second, a CellTiter-Glo^®^ Luminescent Cell Viability Assay was used to quantify ATP levels in treated and untreated cells. A decrease in ATP in treated compared to untreated cells is indicative of a decrease in the number of metabolically active cells (Fig 2B). ATP levels decreased in both ATF3^+/+^ and ATF3^−/−^ MEFs following treatment with IGG and PB, however the surviving fraction of treated ATF3^−/−^ MEFs was significantly greater than the surviving fraction of control MEFs. ATF3^−/−^ MEFs were again protected from the cytotoxic effects of both PB and IGG. Taken together, ATF3 expressing MEFs are more sensitive to splicing inhibitors than their ATF3^−/−^ counterparts, as assessed by DNA fragmentation, membrane integrity and ATP content. ATF3 is important in multiple cell death pathways [7, 11, 14] but its contribution to cell sensitivity to the spliceosome inhibitors was unknown.

### Splicing inhibitors increase ATF3 expression in human cells

Our results provide compelling genetic evidence that ATF3 contributes to cell death induced by spliceosome inhibitors in MEFs. We sought to determine if splicing inhibitors similarly led to ATF3 induction in human cells. Therefore, we measured the expression of ATF3 mRNA and protein in response to treatment with both splicing inhibitors in colorectal carcinoma and cervical carcinoma cells (HCT116 and HeLa cells, respectively). As detected in MEFs, ATF3 mRNA increased significantly in response to PB in both cell lines while the small increase in ATF3 mRNA in IGG treated cells was not statistically significant (Fig. 3A and B). Again, ATF3 protein was induced more in response to PB than IGG (Fig. 3C and D). Both cancer cell lines were exposed to either IGG or PB and 48 hours later apoptosis was assessed by flow cytometric analysis of subG1 DNA content. Both splicing inhibitors resulted in an increase in the fraction of cells with sub G1 DNA content under the same conditions with higher levels of apoptosis detected in response to PB (Figs 3 E and F), consistent with greater fold increases in ATF3 expression. These results support the concept that ATF3 is induced upon spliceosome disruption and that this could contribute to cell death in human cells as well.

**Fig 3.**
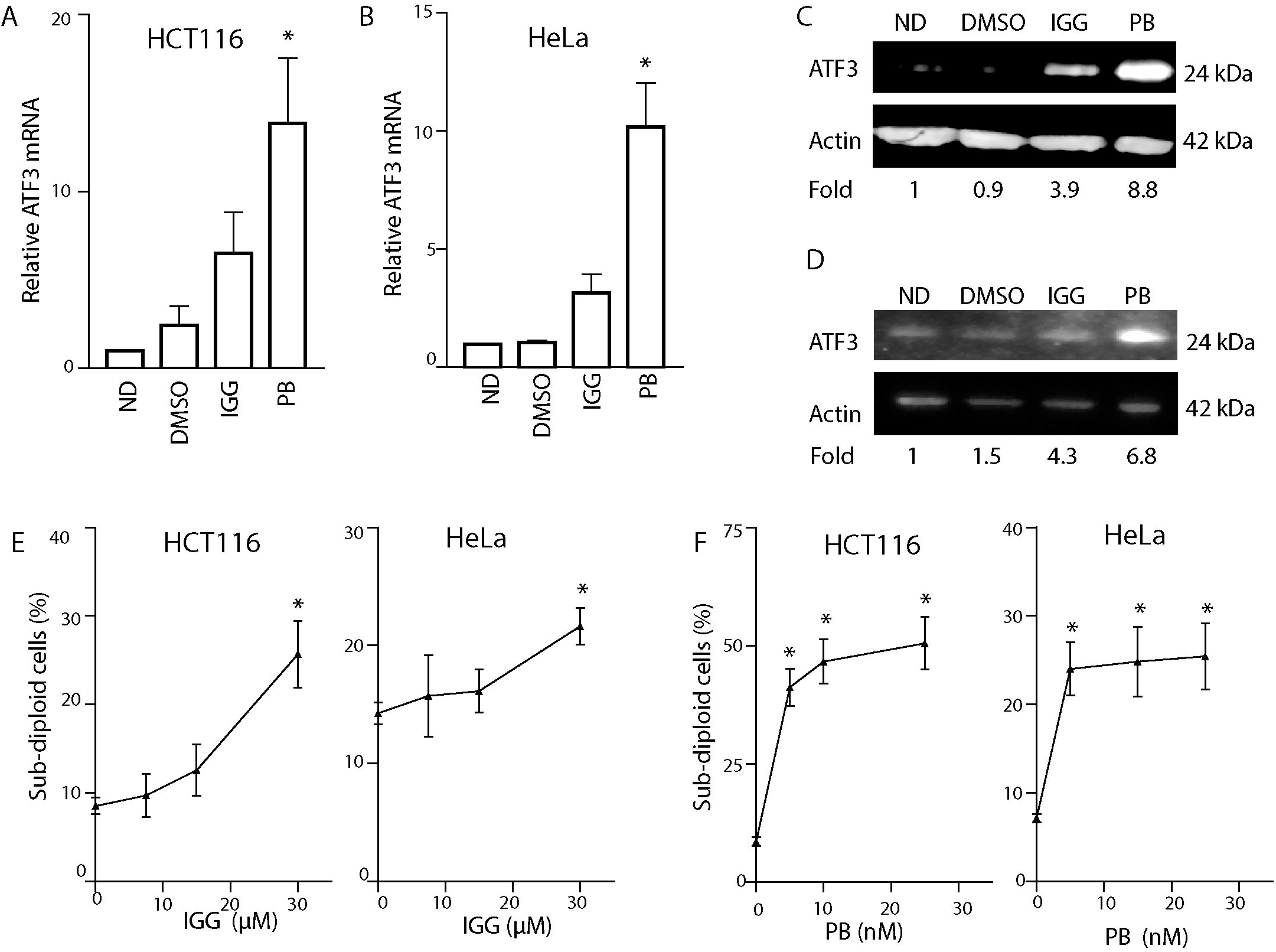
ATF3 expression is increased in human cells exposed to IGG and PB. HCT116 and HeLa cells were left untreated, exposed to DMSO, 30μM IGG or 25nM PB for 24h (A-D) or 48 hours (E and F). (A-D) ATF3 mRNA and protein levels were measured using qRT-PCR and immunoblotting, respectively, in HCT116 and HeLa cells. The mean fold change determined from 3 independent experiments is presented below the representative lane of the immunoblot. (E and F) Flow cytometric analysis of sub G1 DNA content was performed 48 hours following treatment of HCT116 and HeLa cells in response to IGG and PG, as indicated in each panel. Each point represented the mean, +/− SEM from at least three independent experiments. Statistical significance was determined using a one-way ANOVA followed by a Tukey’s post-test (*p<0.05).

### Knockdown of ATF3 protects HeLa cells from spliceosome inhibitors

To determine if ATF3 contributes to cell sensitivity in response to splicing inhibitors in these human cancer cell lines, we performed siRNA mediated knockdown of ATF3 in HeLa cells. The siRNA transfected cells were seeded to smaller dishes to perform all assays from the same transfected sample. RNA interference led to decreased ATF3 expression and prevented the full induction of ATF3 following drug exposure (Figs 4A and B). As expected, ATF3 knockdown protected HeLa cells against the cytotoxic effects of these splicing inhibitors, although only protection against PB was statistically significant (Fig 4C). These results support the concept that ATF3 is induced upon spliceosome disruption and that this contributes to cell death in both murine and human cells.

**Fig 4.**
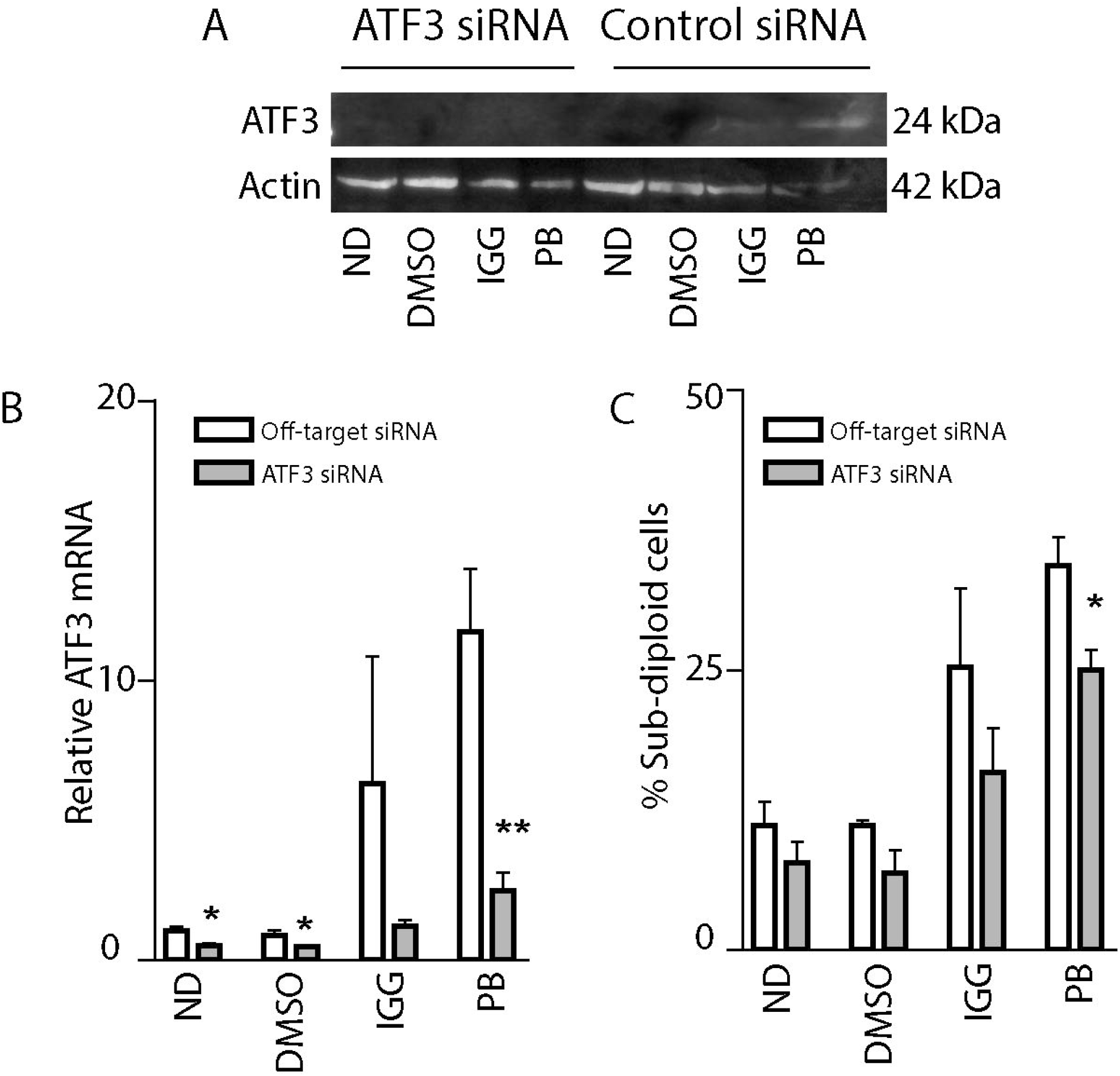
Knockdown of ATF3 protects HeLa cells from IGG-and PB-induced apoptosis. HeLa cells were transfected with control non-targeting or anti-ATF3 siRNAs, split into separate dishes for RNA, protein, and apoptosis analysis. Transfected cells were subsequently exposed to the indicated drugs (30 μM IGG or 25 nM PB), no drug control (ND) or vehicle control (DMSO) for up to 48 hours. (A and B) Twenty-four hours following treatment, protein and RNA samples were collected for immunoblot (A) and qRT-PCR analysis (B). Similar immunoblot results were obtained over from three independent experiments. Each value in B represents the mean +/− SEM determined from three independent experiments. (C) Sensitivity to IGG- and PB-induced apoptosis was assessed by flow cytometric analysis of subG1 DNA content. Each point represents the mean +/− SEM determined from three independent experiments. The * and ** in B and C indicate that the value is significantly different from the corresponding non-targeted siRNA transfected control sample (p<0.05 and p<0.01, respectively), as determined using unpaired t-tests.

## Discussion

The production of functional mRNAs requires the synthesis of pre-mRNA that is coordinated with co-transcriptional mRNA processing steps including pre-mRNA splicing [15–17]. The inhibition of pre-mRNA splicing delays or prevents the production of functional mRNAs and indirectly the encoded proteins. How splicing abnormalities are sensed and transmitted to cell death pathways remains unclear. In the present work, we measured cellular responses to two spliceosome inhibitors, IGG and PB. We found that both drugs led to an increase in ATF3 expression in human colon cancer cells and MEFs. Parental MEFs, expressing ATF3, were sensitive to both inhibitors leading to a large decrease in the viable cell population. However, cell viability was significantly higher in ATF3−/−, as assessed in 3 independent assays. Subtle quantitative differences were detected among the assays that reflect the timing of individual assays, the measured characteristic of cell death (DNA fragmentation, loss of membrane integrity or loss of ATP) and the sensitivity of individual assays. In addition, CellTiter-Glo^®^ Luminescent Cell Viability Assay is not determined on a single cell basis and it is very sensitive [18] so it isn’t entirely surprising that we detected greater apparent cell death with this assay. Despite these differences, all three assays support a model in which ATF3 is a major regulator of mammalian cell sensitivity to these pre-mRNA splicing inhibitors.

To address the importance of ATF3 in determining the sensitivity of human cells to spliceosome inhibitors, we used siRNAs to target ATF3 in HeLa cells. We were able to knockdown ATF3 by about 50% prior to treatment and RNA interference prevented the induction of ATF3 by IGG and PB. Consistent with data obtained in MEFs, we found that reduced ATF3 expression was associated with decreased sensitivity to apoptosis. Therefore, ATF3 appears to contribute to IGG- and PB-induced cell death in human cells, as well. It is noteworthy that PB led to higher

ATF3 levels and greater sensitivity to cell death in all cell lines, consistent with ATF3-dependent apoptosis. This is remarkable in that the concentrations of PB used here are well below the reported IC50 for inhibition of *in vitro* splicing (90 nM) while there is less ATF3 and cell death induced by IGG at its reported in vitro IC50 (30 μM) [4, 19]. It is unclear why we detected differences in ATF3-dependent apoptosis but this adds to a growing list of differences reported in the literature.

The two spliceosome inhibitors tested target distinct pre-spliceosome complexes. PB interacts with the SF3B1 subunit of the U2 snRNP in the complex A [5] whereas IGG acts somewhat later preventing the transition from the A to B complex through an unknown target [4]. Even though both drugs inhibit pre-mRNA splicing at similar stages and induce apoptosis through an ATF3-dependent mechanism, a variety of differences in gene expression and biological effects have been described. For example, we have previously reported that these two drugs can have pronounced effects on cell cycle distribution but these changes were not identical [20]. Two parameter flow cytometric analysis of bromodeoxyuridine-labelled cells indicated that IGG led to multiple cell cycle alterations but the most prominent was to an S phase arrest [20]. In partial contrast, PB-treated cells were blocked predominantly in G_1_ but also exhibited a more modest S phase arrest than IGG-treated cells [20]. Meanwhile, both drugs led to decreased M phase assessed by loss of phospho-H3 immunoreactivity [20]. Therefore, there are similarities and differences in the effects of these compounds on cell cycle responses.

Other groups have reported recently that IGG has other unique effects on transcription and protein trafficking. Boswell and coworkers found that IGG decreases transcription elongation, in addition to its effects on pre-mRNA splicing [21]. Strand-specific RNA-sequencing (RNA-seq) was performed in IGG-treated and control cells and relative read number increased within 5 kb of transcription start sites in IGG-treated cells but not in the presence of an SF3B1 inhibitor, meayamycin [22], suggesting that transcription near the 5’end of genes was slow in the presence of IGG [21]. These authors also detected increased polyA site readthrough in IGG treated cells further suggesting that a fundamental problem in transcription and co-transcriptional processing may underlie some of the unique biological effects of IGG [21]. It is noteworthy that these authors still detected a defect in pre-mRNA splicing using RNA-seq analysis in this and previous papers [21, 23, 24]

Lastly, IGG also appeared to inhibit trafficking of the amyloid precursor protein (APP) from endoplasmic reticulum (ER) to the Golgi [24]and lysosome-mediated turnover [13]. In the former, APP levels increased in response to IGG but not the SF3B1 inhibitor spliceostatin [24]. APP was retained in the ER where the protein was stable and accumulated [24]. In the latter paper [13], IGG led to increased lysosomal stress, increased TFEB protein expression (a regulator of lysosomal biogenesis) and the accumulation of polyubiquitinated proteins under nutrient starving conditions suggesting that IGG may be inhibiting proteasomes and altering lysosomal function under some conditions [13]. Intriguingly, IGG-induced TFEB activated ATF3 indirectly through ATF4, however it was unclear if ATF3 contributed to cell death detected in this manuscript. There is presently no evidence that PB or any other SF3B1 inhibitors similarly affect protein homeostasis. While these additional functions attributed to IGG may contribute to differences detected in cell cycle regulation [20], they don’t appear to strictly alter the ATF3-dependence of cell death induced by IGG and PB. It is likely that IGG and PB lead to ATF3 induction through a common mechanism related to spliceosome inhibition.

## Conclusions

ATF3 is involved in a variety of stress pathways, and it is likely that a disruption in spliceosome formation is leading to activation of ATF3 through an unknown mechanism. The novel findings here provide evidence that splicing inhibitors lead to activation of ATF3, and that ATF3, in turn, is playing a role in determining sensitivity of cells to these compounds. Our future work will investigate the role ATF3 has on cell sensitivity, and work to determine how each of these splicing inhibitors specifically led to ATF3 activation.

## Acknowledgements

We would like to thank Dr. Jim Dimitroulakos (Princess Margaret Research Institute, Toronto, ON) for providing mouse embryonic fibroblasts.

